# Impaired Layer Specific Retinal Vascular Reactivity Among Diabetic Subjects

**DOI:** 10.1101/2020.05.15.097717

**Authors:** Maxwell Singer, Bright S. Ashimatey, Xiao Zhou, Zhongdi Chu, Ruikang K. Wang, Amir H. Kashani

## Abstract

**Purpose:** To investigate layer specific retinal vascular reactivity (RVR) in capillaries of diabetic subjects with no or mild non-proliferative diabetic retinopathy (NPDR).

**Methods:** A previously described nonrebreathing apparatus was used to deliver room air, 5% CO_2_, or 100% O_2_ to 41 controls and 22 diabetic subjects (with mild or no NPDR) while simultaneously acquiring fovea-centered 3×3mm^2^ Swept-Source Optical Coherence Tomography Angiography. Vessel skeleton density (VSD) and vessel diameter index (VDI) were calculated for each gas condition for the superficial retinal layer (SRL) and deep retinal layer (DRL). The superficial layer analysis excluded regions of arterioles and venules. Data analysis was performed using mixed factorial analysis of covariance stratified by diabetic status. All models were adjusted for age, gender, and hypertension.

**Results:** Among controls, there was a significant difference in capillary VSD between all gas conditions (p<0.001). This difference was present in both the SRL and DRL. Among diabetics, there was no significant difference in response to CO_2_ conditions in the SRL (p=0.072), and a blunted response to both CO_2_ and O_2_ in the DRL. A significant gas effect was detected in the capillary VDI in the SRL of controls (p=0.001), which was driven by higher VDI in the oxygen condition compared to that of carbon dioxide.

**Conclusions:** Impairment in RVR in diabetic subjects is driven largely by a decrease in the magnitude of the capillary response to O_2_ in the DRL as well as almost complete attenuation of capillary CO_2_ response in all layers. These layer and gas specific impairments in diabetics seem to occur early in the disease and to be driven primarily at the capillary level.

## Introduction

Blood flow through the retinal vasculature changes according to the metabolic needs of local tissue and the presence or absence of metabolites in the local environment.(1) One example of this spatial and temporal heterogeneity in blood flow is the reactivity of retinal vessels to blood gas perturbations. Retinal vessels change in caliber, dilating in hypercapnic conditions and constricting in hyperoxic conditions.(2–7) Such changes are mediated by regulatory mechanisms that allow for decentralized regulation of blood flow within the larger vessels of the retina.(8) For example, retinal arteries and arterioles can change in diameter in response to changes in metabolic demand or supply.(9–11)

This retinal vascular reactivity (RVR) has been shown to be blunted in diabetics (12–16) even in the absence of clinically apparent diabetic retinopathy (DR).(2) Most studies have demonstrated reactivity changes in arterioles or venules, but few have demonstrated impairment at the capillary level.(1,6) Unlike many other imaging modalities that have been used to assess retinal vascular reactivity at the capillary level, Optical Coherence Tomography Angiography (OCTA) is an FDA approved technology for commercially use.(17,18) Our group has recently used Swept-Source OCTA (SS-OCTA) to investigate the impact of room air, hypercapnia and hyperoxia conditions on retinal capillaries in full thickness retina between controls and subjects with diabetic retinopathy. That study concluded that subjects with DR preferentially responded to hyperoxia but not hypercapnia, while the healthy controls responded to both.(19)

There is currently growing evidence that the deep retinal vascular plexus is more severely damaged by diabetes than the superficial plexus, with some studies showing that even in the absence of clinically detectible retinopathy, the deep vascular plexus shows lower capillary density than in controls.(20–24) Given that vascular damage may occur early in disease and vary by retinal layer, we expand our previous studies by investigating layer specific retinal vascular reactivity between healthy controls and subjects with mild non-proliferative diabetic retinopathy (NPDR) or no DR.

## Methods

This was a prospective, observational study of diabetic patients and healthy controls. This study was approved by the institutional review board of the University of Southern California (USC) and adhered to the tenets of the Declaration of Helsinki.

### Subjects

Subjects were recruited at the USC Roski Eye Institute and informed consents were obtained from each subject. Subjects were excluded if they had any past ocular history of glaucoma or retinal surgery, or past medical history of vascular diseases including heart disease, surgery, or vasculitis. Only those diabetic subjects without clinically apparent DR or with mild NPDR were included. All subjects with significant media opacity such as cataracts or any other obscuration of the fundus were excluded. Subjects who reported a history of syncope, shortness of breath, lung disease, congestive heart failure, or recent hospitalization of greater than 1-day duration within the last 12 months were also excluded to minimize risk of adverse events during gas breathing. All eyes were imaged at the USC Roski Eye Institute between October 2017 and February 2020. Approximately, 46% of controls and 36% of diabetics overlapped with those from our previous study by Ashimatey et al.(19)

### Gas Delivery System

In order to create conditions of physiological hyperoxia and hypercarbia, a nonrebreathing apparatus was used to deliver gas mixtures. This gas delivery system has been described by Ashimatey et al.,(19) and a step by step visual-illustration of the procedure is demonstrated in Kushner-Lenhoff et al.(25) In brief, three gas mixtures were delivered through a non-rebreathing gas apparatus to the subjects in the following order during the same visit: room air, a mixture of 5% carbon dioxide, atmospheric oxygen and balanced nitrogen, and finally 100% oxygen. There was a 10-minute refractory period between the hypercapnic and hyperoxic conditions to allow washout of the previous gas condition. During delivery of the gases, a fingertip pulse oximeter was used to monitor oxygen saturation. Testing was stopped if the oxygen saturation fell below 94% for any subject at any time.

### SS-OCTA Imaging Protocol

During gas breathing, images were acquired using a 100kHz SS-OCTA system (PLEX Elite 9000, software version 1.7; Carl Zeiss Meditec, Dublin, CA, USA). The light source has a central wavelength of 1060 nm and a bandwidth of 100 nm, providing an axial resolution of ∼ 6μm and lateral resolution of ∼ 20μm (estimated at the surface of retina). The device uses an optical microangiopathy (OMAG) algorithm(26,27) to determine the motion signal, which is interpreted as representing the flow of erythrocytes. Three OCTA volumes (3×3mm field-of-view) centered on the fovea were acquired starting 60 seconds after the onset of each gas breathing condition. Each OCTA volume is formed by 300 horizontal scans (B-scans) with each B-scan formed by 300 A-scans of 1536 pixels. One eye per patient was imaged, based on good fixation ability, absence of co-morbid disease and media opacity. If both eyes were suitable, the imaged eye was chosen randomly. Images with poor decentration of the foveal avascular zone (FAZ), signal strength less than seven, significant motion artifacts or vitreous floaters were excluded.

One representative image, the image with the highest signal strength and fewest artifacts, was selected per condition. The volumes were then automatically segmented into *en face* superficial retinal layer (SRL) and deep retinal layer (DRL). The segmentation software used is commercially available on the OCTA device and has been previously described by our group in the past.(28,29) The SRL extends from the inner limiting membrane (ILM) down through the inner plexiform layer (IPL), which is approximated as 70% of the thickness between the ILM and outer plexiform layer (OPL). The DRL extends from the IPL to the OPL, which is estimated to be 110μm above the retinal pigment epithelium (RPE). The *en face* DRL for the best images were projection resolved using the commercially available projection artifact removal feature on the device.(30,31)

### Quantitative Vessel Morphometric Measures

Quantitative analysis with custom MATLAB software was performed on SRL and DRL images to calculate measures that have been described previously.(32) Vessel area density (VAD) represents the total area of the image occupied by vessels and is calculated by dividing the area occupied by vessel pixels by the dimensions of the image after image binarization. Vessel skeleton density (VSD) represents the total length of the blood vessels. VSD is calculated by skeletonizing the diameter of each vessel to one pixel and then dividing the pixels that represent all the skeletonized vessels by the total area of the image. Vessel diameter index represents the caliber of the blood vessels and is calculated by dividing VAD by VSD.(28,33,34)

The central 3×3mm foveal centered region of the macula contains small arterioles which may preferentially drive the RVR response. In order to examine the isolated effect of gas conditions on capillaries only, a custom MATLAB software was developed specifically to exclude large vessels that occur in the SRL. The software uses Hessian filter and pixel intensity values to detect large vessels and exclude them from capillary analysis (Figure 1). VSD is then calculated by dividing the remaining skeletonized capillaries by all the pixels of the image that were not occupied by large vessels. In addition, VSD, VDI, and VAD of the SRL with the large vessels included was also quantified for comparison. Unless otherwise specified, results presented are those with large vessels excluded.

**Fig 1.**
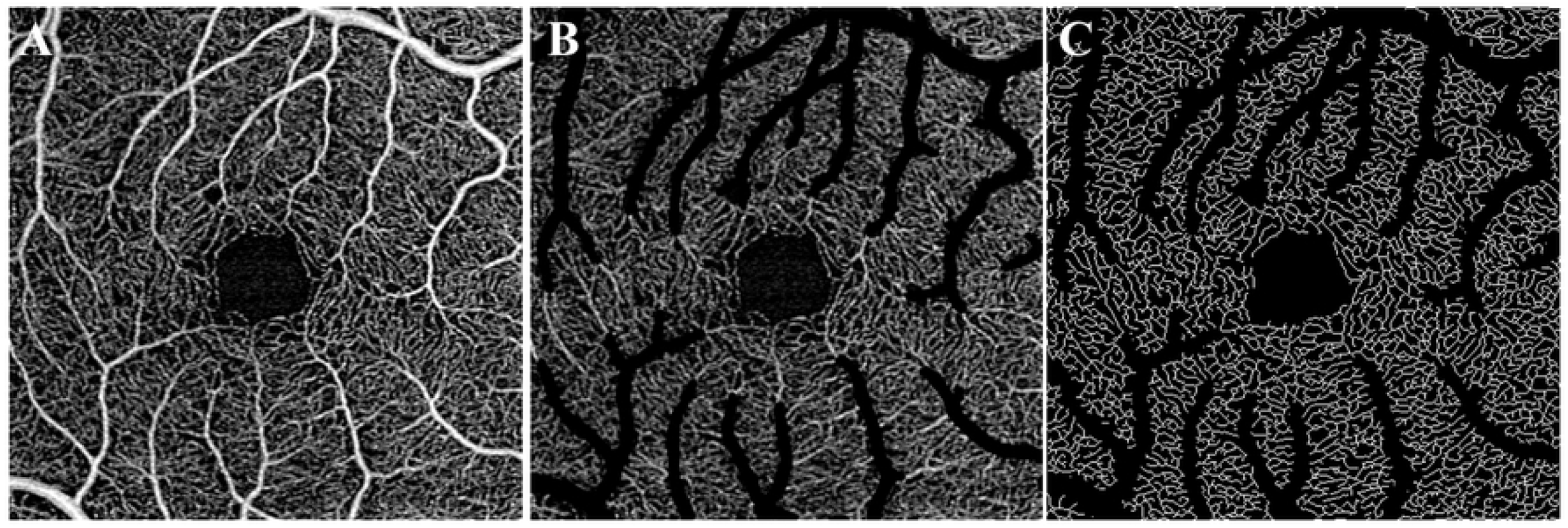
Large vessel exclusion of en face superficial retinal layer (SRL) 3×3mm SS-OCTA of control subject acquired during room air condition. (A) original image (B) original image with large vessels removed (C) image after large vessels excluded and remaining capillaries skeletonized

### Statistical Analysis

A 2 (layers: DRL, SRL) x 3 (gas conditions: room air, oxygen, carbon dioxide) mixed factorial analysis of covariance (ANCOVA) model with repeated effects of layer and breathing condition was used to assess differences in the OCTA metrics (VSD, VAD and VDI) by layer, gas breathing condition, and their interaction. Due to limitations in sample size and power to detect a three-way layer-gas-diabetes interaction, separate models were run for diabetic versus normal patients and for VSD measurements obtained for the images. All models were adjusted for age, gender, and hypertension. Differences of the differences in VSD by gas breathing condition between layers were estimated from the models. All statistical tests were two-sided with an alpha level set at 0.05 for statistical significance. The Bonferroni method was applied for multiple comparisons. All statistical analyses were performed using SPSS version 25 (IBM SPSS Statistics for Windows, Version 25.0. Armonk, NY).

## Results

Our study included 41 control (mean age 53.0 ± 18.9) and 22 diabetic subjects (mean age 53.7 ± 16.7). Table 1 summarizes the subject demographics. Figure 2 shows qualitative representations of the decorrelation signal intensity and capillary density under different gas breathing conditions in the SRL and DRL for a control subject. The same representations are shown for a diabetic subject in Supplementary Figure S1. Subtle signal intensity and the apparent density of capillaries in the control subject were qualitatively increased in the hypercapnic condition compared to the other two conditions. Quantitative analysis confirmed this finding by showing that significant effects on VSD of gas breathing condition, layer, and the gas-layer interactions were present for both controls and diabetics, after controlling for age, gender, and hypertension (all p<0.001). Below we elaborate on these findings in more detail.

**TABLE 1.** Subject Demographics.

|  | Disease Classification |  |
| --- | --- | --- |
|  | Controls | Diabetics |
| Number (Subjects) | 41 | 22 |
| Age | 53.0 ± 18.9 | 53.7 ± 16.7 |
| Female gender | 20 (48.8%) | 9 (40.9%) |
| Hypertension | 10 (24.4%) | 6 (27.3%) |
| Severity of DR | N/A | 15 no DR,<br>7 mild NPDR |

The number, age, gender, hypertension status, and ocular diagnosis of subjects is summarized.

**Fig 2.**
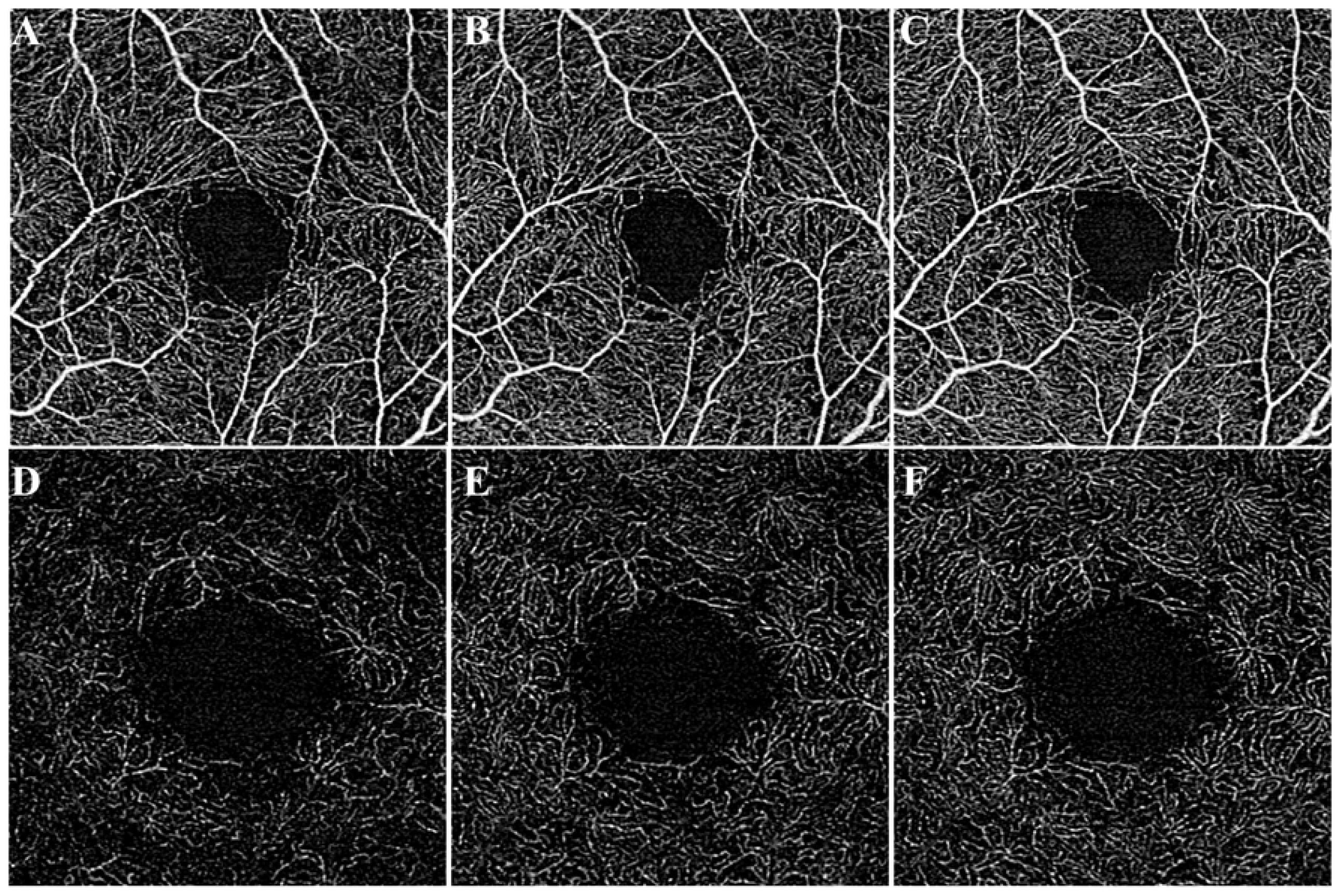
SS-OCTA of control subject during different gas conditions. (A-C) 3×3mm en face superficial retinal layer image of blood flow around central macula during oxygen condition (A), room air condition (B), and carbon dioxide condition (C). (D-E) 3×3mm en face deep retinal layer image of blood flow around central macula during oxygen condition (D), room air condition (E), and carbon dioxide condition (F). A qualitative increase in signal was apparent in the hypercapnic condition when compared to the other two conditions.

### Retinal vascular reactivity to CO_2_ is absent in SRL of diabetic subjects

There was no significant difference in mean VSD of diabetic subjects for the pairwise comparisons between room air and CO_2_ gas conditions (SRL_RA_ 0.148 ± 0.002 vs SRL_CO2_ 0.146 ± 0.002, p=0.072). As Figure 3 shows, the reactivity of retinal capillaries to CO_2_ was not only absent in the SRL of diabetic subjects but the mean VSD under CO_2_ was paradoxically lower than RA. In contrast, in the SRL of control subjects, there was a robust difference in all pairwise comparisons of the mean VSD among the three gas conditions in the expected directions (SRL_RA_ 0.154 ± 0.001 vs SRL_CO2_ 0.156 ± 0.002 vs SRL_O2_ 0.150 ± 0.002, p<0.017). Results for VAD were consistent with those of VSD, and the VSD results were unchanged when large vessels were not excluded from the SRL (see S2 Figure and S3 Figure in Supplement). The omnibus findings of the mixed factorial analysis confirmed a significant gas condition (p<0.001), layer (p<0.001) and gas-layer interaction (p<0.001) in the controls. In diabetic subjects, the mixed factorial analysis confirmed that there was a significant effect of gas condition (p=0.001), mainly driven by the response to O_2_, layer (p<0.001) as well as a significant gas-layer effect (p=0.004).

**Fig 3.**
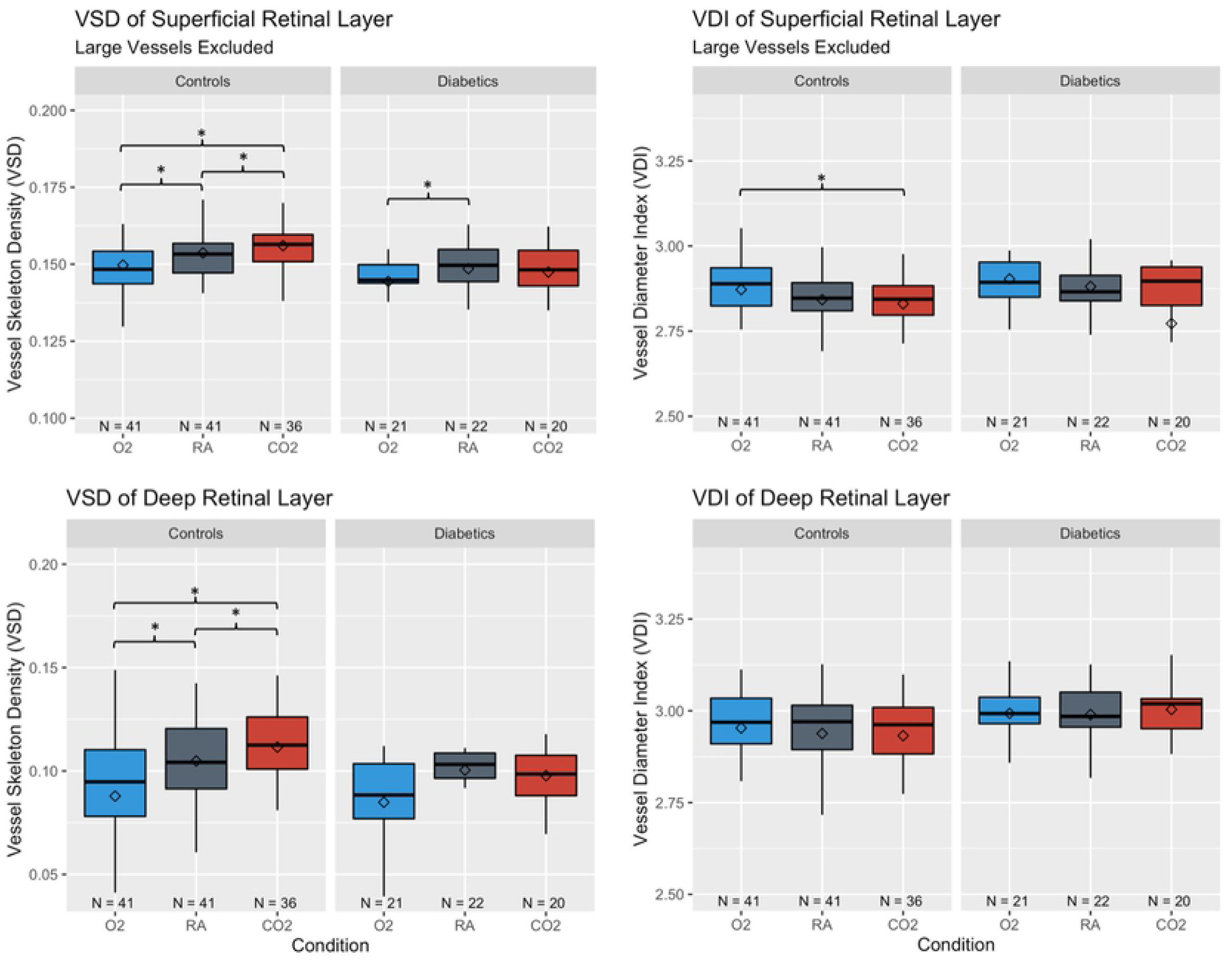
Vessel skeleton density (VSD) and vessel diameter index (VDI) of control and diabetic subjects under gas conditions. *Whiskers* indicate highest or lowest point within 1.5 times the interquartile range from the upper or lower quartile. *Diamonds* indicate the mean. *Stars* indicate significance for pairwise comparison between conditions from the ANCOVA model based on Bonferroni adjusted p-value of 0.017.

For the VDI metrics, there was a significant gas effect in the SRL of control subjects (SRL_RA_ 2.84 ± 0.01 vs SRL_CO2_ 2.83 ± 0.01 vs SRL_O2_ 2.87 ± 0.02, p=0.001) which was driven primarily by the difference in VDI between O_2_ and CO_2_ (p=0.002). Gas effect on the VDI metric in the SRL was not present in diabetic subjects (SRL_RA_ 2.88 ± 0.03 vs SRL_CO2_ 2.77 ± 0.03 vs SRL_O2_ 2.90 ± 0.02, p=0.44). In the controls, the mixed factorial analysis confirmed that there was a significant effect of both gas conditions (p=0.005) and layer (p<0.001) on VDI but no significant gas-layer interaction (p=0.29). In diabetic subjects, the mixed factorial analysis neither confirmed a significant effect of gas condition (p=0.51) nor a significant gas-layer effect (p=0.38) but did show significant differences in between layers (p=0.009). The VDI results were similar when large vessels were not excluded from the SRL (see S3 Figure).

### Retinal vascular reactivity to CO_2_ is absent in DRL of diabetic subjects

Unlike the controls with significant pairwise comparisons between gas conditions (p<0.017), all pairwise comparisons of VSD between gas conditions in the DRL of diabetic subjects were non-significant. The mixed factorial analysis of the gas effect in the diabetics however showed significance (p=0.002) and appeared to be driven by the response to hyperoxia. Notably, as in the SRL, the difference in the VSD of the DRL under hypercapnic conditions was in a paradoxical direction and reflects an aberrant physiological response in the diabetic subjects. In the controls the mixed factorial analysis confirmed significant gas effects on VSD in the DRL (DRL_RA_ 0.105 ± 0.003, DRL_CO2_ 0.112 ± 0.003, DRL_O2_ 0.087 ± 0.006, p<0.001). Results for VAD in the DRL were consistent with those of VSD (see S2 Figure). Neither the controls nor subjects with diabetes had a significant gas effect on the VDI within the DRL.

## Discussion

We investigated layer specific RVR during physiologically salient changes in inhaled oxygen and carbon dioxide in diabetic subjects and non-diabetic controls using SS-OCTA. Consistent with our previous findings (19) we found retinal vascular reactivity impairment in diabetic subjects can be assessed with OCTA and show that even in the early stages of the disease the impairment to hypercapnia previously reported are present. Whereas our earlier study consisted entirely of DR subjects with the full range of disease severity, our current cohort of diabetic subjects included only those without DR (68%) or those with mild NPDR. We extend our previous findings by demonstrating that the impairment in RVR in diabetic subjects is present in both the SRL and DRL and is more profound in the DRL. Namely, there was no significant effect of any gas condition on apparent capillary density in the DRL among diabetic subjects by pairwise comparison. Lastly, we show that the RVR findings are present even when larger arterioles and venules are excluded from the images. This suggests that capillaries themselves mediate the RVR and that this physiological response is abnormal in diabetic retinopathy.

We found that retinal vascular reactivity to hyperoxia was preserved in the SRL of diabetic subjects but not significant in the DRL, even in the early stages of the disease. This is consistent with the recent report by Meshi et. al. that among diabetic subjects without DR, capillary density in the deep capillary plexus but not the superficial capillary plexus was significantly lower than that of controls.(24) Various other studies have reported a similar result for type 1 diabetics with no or mild DR.(21–23) It may therefore be that capillary damage in the DRL, including the ability to autoregulate, is more extensive than damage to the SRL in early disease.

We primarily used a measure of capillary density, VSD, which represents the length of capillaries in the image. Changes to VSD with the gas conditions could possibly show perfusion of previously unperfused vessels or vice versa, but more likely reflects functional changes in flow through capillary segments. Due to the detection sensitivity limitations of the OCTA imaging in which flow below a certain threshold cannot be resolved, loss of signal for a given capillary segment between gas conditions may be from decreased perfusion through that segment outside the detection limits of the device. Differences in the VSD measures under the different gas conditions are thus convincing evidence of the change in perfusion per unit area, apart from changes in the vessel caliber induced by the gas conditions, and with minimal masking from the capillary diameter. Since other measures such as capillary diameter are an essential component of the vessel reactivity, we also included measures of VDI and VAD, which both capture the vessel diameter. VAD showed consistent findings with our VSD results.

VDI results were systematically lower with increased CO_2_ content in the controls. As VDI is a ratio of total vessel area (VAD) divided by skeletonized vessel length (VSD), the low VDI observed in the carbon dioxide condition compared to the oxygen condition is likely driven by a relative increase in vessel length compared to vessel width. We interpreted this to mean that the changes in the vessel area density assessible with OCTA under the gas perturbations conditions are largely driven by the absence or presence of flow, rather than changes to flow caliber at the capillary level. Furthermore, taking into account how our VDI metric was computed, a non-significant change in VDI (such as in the DRL of controls) is not necessarily equivalent to non-dilation of the vessels, but could potentially mean that any vessel dilation is neutralized by an increase in appearance of new vessel segments.

The findings of our study come with some limitations which we have tried to mitigate. Namely our sample size was small. Even though our sample size was small our data was representative of the general trends previously described by our group and others, namely, that the mean VSD for the diabetic subjects was lower than the controls and VSD in the SRL was consistently greater than in the DRL. Notably, over half of our cohort of diabetic subjects did not have retinopathy and the remainder had mild NPDR. The current results seem then to reinforce the notion that changes in RVR are an early event in diabetes and warrant further investigation as a preclinical biomarker of disease severity.

One other limitation of our study is that the gas conditions we used were relatively mild to ensure subject comfort and safety. Nevertheless, the methodology is very useful because it can be applied using commercially available devices and on living human subjects with reasonable and reliable effect sizes. We have provided a detailed description of the methodology and its strengths and limitations in our previous publication Ashimatey et al.(19) as well as in Kushner-Lenhoff et al.(25)

To conclude, we have expanded upon our earlier study assessing retinal vascular reactivity in response to gas perturbation to investigate layer specific changes in diabetic subjects without DR or with mild NPDR and to localize those changes to capillaries rather than larger vessels. We found loss of reactivity to hypercapnia in both the superficial and deep retinal plexuses with preserved reactivity to hyperoxia in the superficial layer. These findings specify layer specific changes to retinal vascular reactivity that may play a useful role in disease detection before the onset of clinically detectable diabetic retinopathy and expand our understanding of the pathophysiology of DR.

## Acknowledgments

The authors would like to thank Melissa Mert and Luis de Sisternes for providing statistical and OCTA-related technical advice respectively during the manuscript development.

## Supplement

**S1 Figure.**
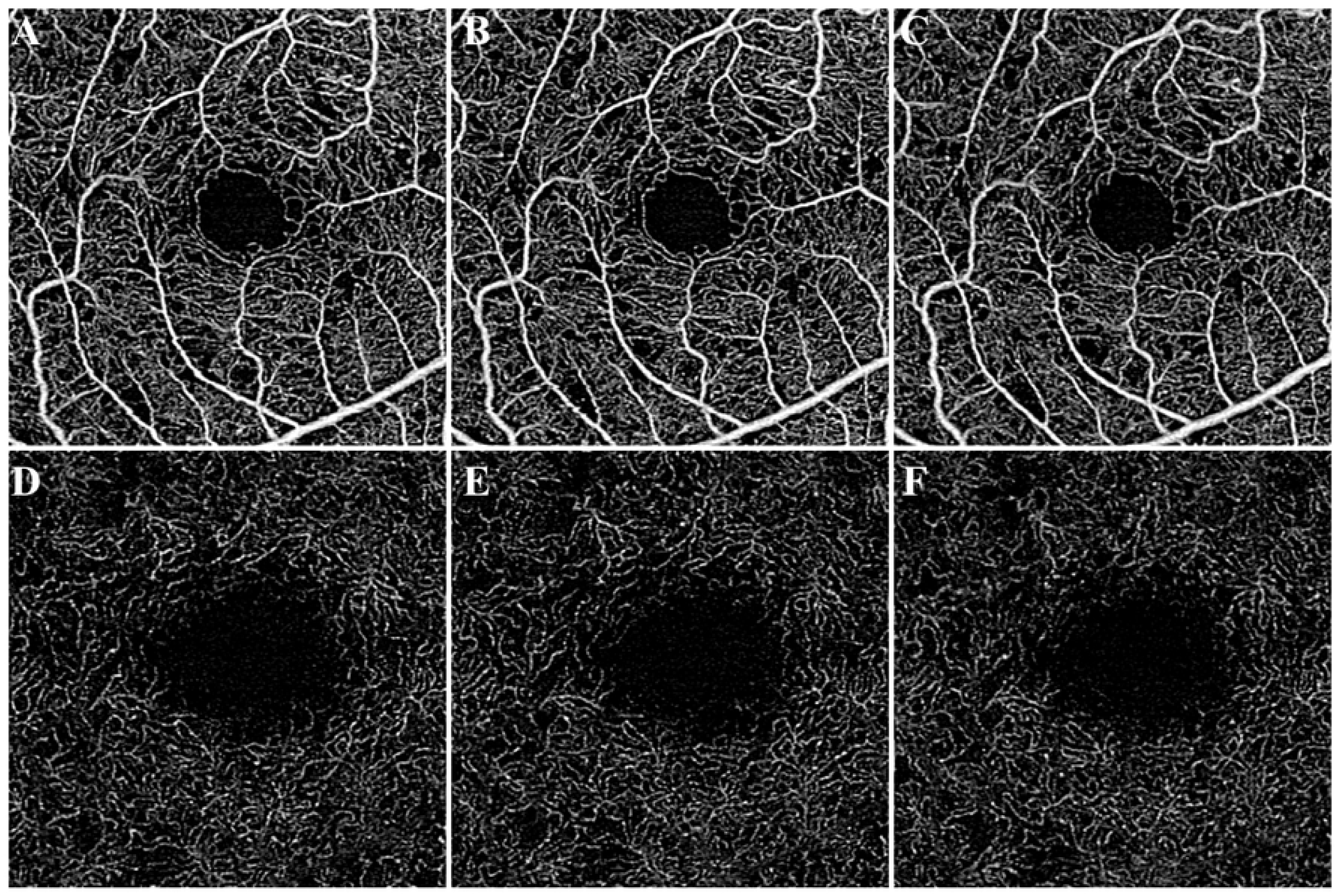
SS-OCTA of diabetic subject without diabetic retinopathy during different gas conditions. (A-C) 3×3mm en face superficial retinal layer image of blood flow around central macula during oxygen condition (A), room air condition (B), and carbon dioxide condition (C). (D-E) 3×3mm en face deep retinal layer image of blood flow around central macula during oxygen condition (D), room air condition (E), and carbon dioxide condition (F).

**S2 Figure.**
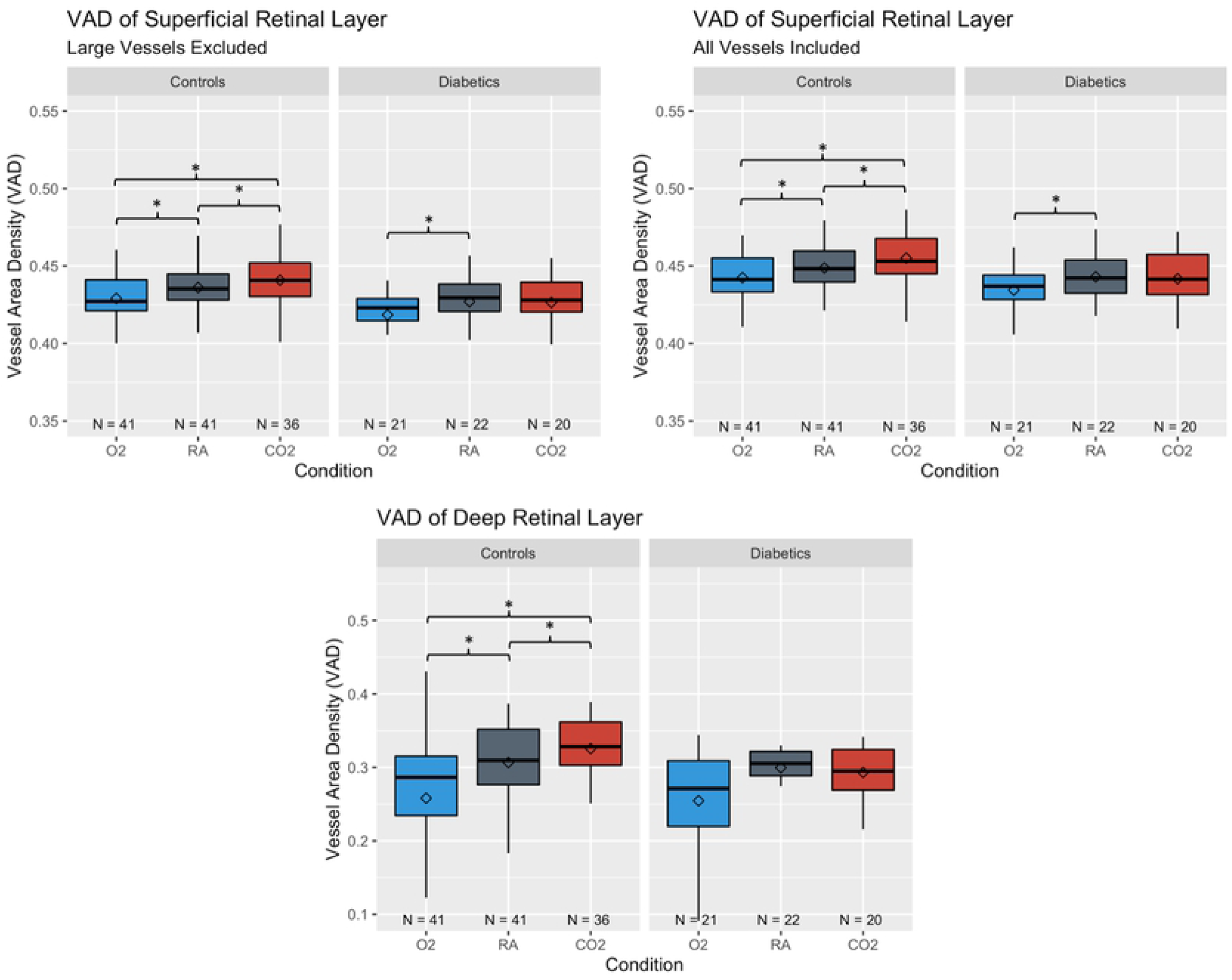
Vessel area density (VAD) of control and diabetic subjects under gas conditions. *Whiskers* indicate highest or lowest point within 1.5 times the interquartile range from the upper or lower quartile. *Diamonds* indicate the mean. *Stars* indicate significance for pairwise comparison between conditions from the ANCOVA model based on Bonferroni adjusted p-value of 0.017.

**S3 Figure.**
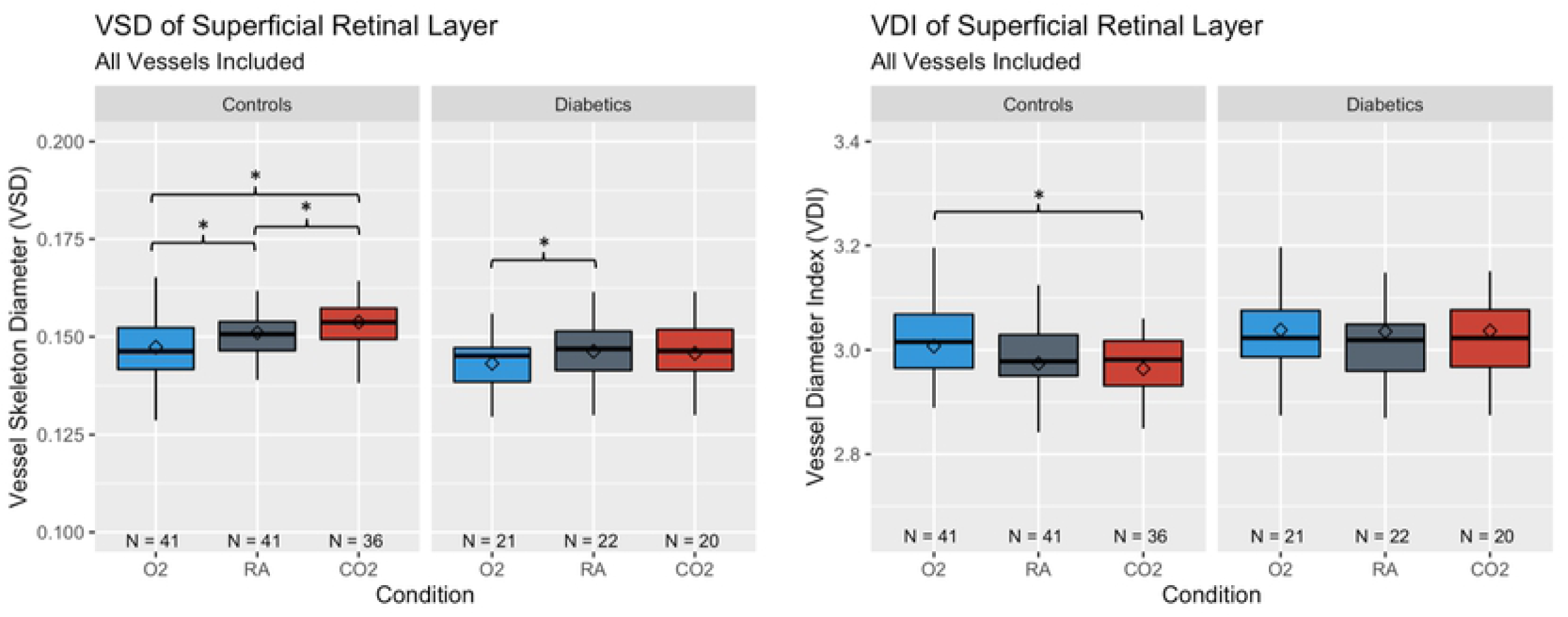
Vessel skeleton density (VSD) and vessel diameter index (VDI) of control and diabetic subjects under gas conditions using all vessel analysis in the superficial retinal layer. *Whiskers* indicate highest or lowest point within 1.5 times the interquartile range from the upper or lower quartile. *Diamonds* indicate the mean. *Stars* indicate significance for pairwise comparison between conditions from the ANCOVA model based on Bonferroni adjusted p-value of 0.017.

